# Landscape drivers and effectiveness of pest control by insectivorous birds in a landscape-dominant woody crop

**DOI:** 10.1101/2020.03.07.981845

**Authors:** Carlos Martínez-Núñez, Pedro J. Rey, Antonio J. Manzaneda, Daniel García, Rubén Tarifa, José L. Molina, Teresa Salido

## Abstract

Avian-mediated pest control is a significant ecosystem service with important economic implications. However, there is an overall paucity of experimental information about how landscape simplification affect its current level. Information on pest control by birds is missing in some permanent agroecosystems of worldwide importance, like olive orchards, that dominate vast areas in the Mediterranean region.

We assess the effectiveness of insectivorous birds for controlling the two main pest insects in olive orchards and explore the effects of landscape complexity and distance to semi-natural patches on avian insectivore abundance and pest control. For this, we combine bird surveys with field experiments (branch exclusions and pest plasticine models) at the regional scale.

Landscape heterogeneity increased the abundance and richness of insectivorous birds, which were also more abundant and diverse in semi-natural patches, compared to the farm olive matrix. Experiments evidenced that pest control by birds (measured as attack rates to plasticine models and pest damage) in the studied olive orchards is negligible, while pests were overall abundant and pest damage was high on most farms. This raises alarms about the status of avian pest control in this agroecosystem.

Although landscape heterogeneity increased the abundance/richness of insectivorous birds, and favored some forest species, insectivorous bird abundance seems diluted in relation to prey availability in all landscapes. Thus, pest control by birds seems currently unsuccessful in olive orchards. Our results might be evidencing the loss of an ecosystem service due to a generalized massive decline of common and forest insectivorous birds.

**Key message:**

- Olive orchards dominate extensive areas causing important landscape simplification.
- Insectivorous birds are more abundant in semi-natural patches within olive farms.
- Field experiments show a low impact of birds on olive pests and damage.
- Avian-mediated pest biocontrol seems diluted by limited suitable habitat for birds.
- Agri-environmental measures should focus on increasing landscape complexity.

## 1. Introduction

Agricultural intensification is often characterized by monocultures that spread over extensive areas, creating highly homogeneous landscapes (He et al., 2019). Monocultures that dominate vast areas are especially prone to suffer from pest damage (He et al., 2019; Redlich, Martin, & Steffan-Dewenter, 2018), because spatial and biotic homogenization favors the spread and success of specialist pests and also removes natural habitats that host potential natural enemies (Grab, Danforth, Poveda, & Loeb, 2018; Rusch et al., 2016). In this sense, only natural enemies that tolerate the micro-environmental conditions of the culture, or that span/forage in large areas, can be effective pest control agents in landscape-dominant cultures. In woody agroecosystems, insectivorous birds can play an important role controlling pest species because the vertical structure of the culture can be alike their natural habitats, they can forage over wide areas and their high metabolic rate forces them to ingest a considerable amount of energy (preys) (e.g. Bael et al., 2008; García, Miñarro & Martínez-Sastre, 2018; Mols & Visser, 2007). Consequently, insectivorous birds are important biological agents, proven successful for pest control in some agroecosystems (García, Miñarro & Martínez-Sastre, 2018; Mols & Visser, 2007)et al., 2018; Karp et al., 2013). However, there are still numerous agroecosystems of worldwide relevance where the effectiveness of avian mediated pest control and the impact of landscape drivers on it, are unknown (Boesing, Nichols, & Metzger, 2017).

Some landscape components can benefit insectivorous bird communities and mediate their contribution to pest control. For instance, semi-natural patches commonly enhance bird communities and pest control on agroecosystems (Escobar-Ramírez, Grass, Armbrecht, & Tscharntke, 2019). Also, larger natural habitat fragments often favor higher predation pressure by birds (Karp et al., 2013), being this pressure usually reduced with distance to the semi-natural patch (Henri et al., 2015; Jordani, Hasui, & da Silva, 2015). Just recently, some studies have started to experimentally approach landscape effects on this ecosystem service. For instance, Maas, Clough, & Tscharntke (2013) used exclosures to demonstrate how agroforest in cacao plantations increased yields through pest biocontrol by birds and bats. García, Miñarro, & Martínez-Sastre (2018), combining experimental insect pest increase and bird exclosures, revealed that forest insectivorous birds, favored by semi-natural woody vegetation around the farms, can control pest outbreaks and diminish damage in apple orchards. Similarly, Koh (2008) used exclusions to show how birds (associated to natural habitat) protected oil palms from herbivores. Other authors have used plasticine models (commonly used to compare attack rates across gradients) to comparatively approach levels of avian insectivory on pests (reviewed in Bateman, Fleming, & Wolfe, 2017). This method have proven useful to explore the effects of agricultural management and landscape heterogeneity on overall pest control (Rusch, Delbac, Thiéry, & Thi Ery, 2017), avian pest control in vineyards (Barbaro et al., 2017) or even to find global latitudinal/altitudinal patterns on predation rates (Roslin et al., 2017). Also, the abundance of insectivorous birds and their predation on plasticine caterpillars has positively been linked to structural heterogeneity (Bereczki, Ódor, Csóka, Mag, & Báldi, 2014).

Although there is increasing evidence that landscape complexity and semi-natural areas benefit insectivorous birds, still very little mechanistic evidence exists about the landscape drivers affecting insectivorous birds and the top-down cascade implications from insectivores (and other natural enemies) to crop damage. Sometimes, system or species/case-specific effects, such as intraguild predation, lack of effective natural enemies, or functional dilution of the insectivory pressure (due to the low numbers of birds compared to the availability of pests) can lead to low or null impact of natural habitat fragments on pest control (Lewis et al., 2013; Tscharntke et al., 2016). In addition, natural vegetation and landscape complexity can also provide disservices to agroecosystems (e.g. sheltering pests or fruit consumers), but landscape complexity seems to be overall positive, even when accounting for the ecosystem disservices (Gonthier et al., 2019; Linden et al., 2019).

Despite that olive orchards are within the most important agroecosystems worldwide (http://www.fao.org/faostat) and their major insect pests are well-known, we lack experimental information about bird pest control potential in this agroecosystem, and about how it is affected by landscape components. Some studies have addressed the importance of natural vegetation (Paredes et al., 2019) and extensive ground cover management (Paredes et al., 2015; Villa et al., 2016) for olive pest control. However, most studies in this agroecosystem have focused on arthropod natural enemies (Álvarez et al., 2019; Paredes et al. 2015), which seem to play an important role, with important economic implications (Paredes et al., 2019). Elucidating the significance of biocontrol by birds and how it is affected by landscape is especially necessary under the current scenario of progressive intensification that suffers this agroecosystem (Infante-Amate et al., 2016) and the remarked role of olive groves for the conservation of bird biodiversity in the Mediterranean region (Rey, 1993, 2011; Rey et al., 2019). In this study, we conduct field surveys together with two experiments involving plasticine dummies and bird exclosures at the regional scale. We experimentally investigate, for the first time, the predation potential of insectivorous birds on the two main olive tree pest species, the olive moth *Prays oleae* (Bernard,1788) and olive fly *Bactrocera oleae* (Rossi 1790), and the damages they produce. We also test the effects of semi-natural patches attributes (e.g. patch size and distance to the patch) and landscape compositional/configurational heterogeneity on: insectivorous bird richness and abundance (with particular attention to forest insectivores), pest abundance, attack rates on plasticine models, and observed crop damage. Our study is conducted exclusively in organic olive farms to avoid potential confounding effects caused by pesticides. Given the evidence of pest control by avian insectivores in other agroecosystems, and provided the known sensibility of insectivorous birds to landscape heterogeneity and presence of semi-natural habitat patches, we hypothesize that insectivorous birds will exert effective pest control of major insect pests in olive orchards, but their effectiveness would be strongly limited in homogeneous landscapes. Our associated predictions are: (1) olive farms located in heterogeneous landscapes and non-crop plots inside farms, will host a higher abundance/richness of insectivorous birds and especially forest insectivores, as well as particular assemblages characterized by a higher frequency of forest species; (2) bird-excluded branches will suffer more damage by pests than controls (free access to birds) probing that birds are important for pest control, and being this effect stronger in trees adjacent to semi-natural patches; (3) attack rates to pest plasticine models, will be higher in heterogeneous landscapes, and close to semi-natural patches. Alternatively, functional dilution of bird insectivory, due to the vast extent of olive orchards in Andalusia, could be responsible for a limited pest control by birds in the olive-dominated landscapes of the region.

## 2. Materials and methods

### 2.1 Study area and study system

Our study is set in Andalucía (south Spain), the region with the highest density of olive trees worldwide. All surveys and experiments of this study were conducted in nine olive farms that span 311 km distance from east to west (from 5□53’46’’W to 2□64’87’’W) and 190 km distance from north to south (from 38□40’05’’N to 36□78’36 N). They are located across a wide range of landscape complexity and were all organic, thereby not agricultural management effect is here considered. For a map showing these localities and more details, see Martínez□Núñez et al. (2019).

In this work we focus on two pest species: *P. oleae* (the olive moth) and *B. oleae* (the olive fruit fly). The olive moth has a complex life cycle, completely adapted to the phenology of the olive tree. This species has three generations. The phylophagous (leaf generation), a small larva that feeds on olive tree leaves and builds galleries during winter-spring. The anthophagous (flower generation), that feeds on floral buttons and flowers, matches the flowering period of olive trees in Spring. The third one is the carpophagous larva (fruit generation), the most harmful to production, causing premature fruit fall in Autumn (Pelekassis, 1962). The olive fruit fly is an obligated pest of olive trees. The adult flies oviposit their eggs in the fruit, where the larvae develop feeding on the fruit. The tunnels produced by the larvae cause necrosis and possible fruit falling from the tree. In Autumn, the larvae pupae in the olive fruit or in the soil, where they spend the winter (Daane & Johnson, 2010).

### 2.2 Landscape complexity

We calculated several compositional and configurational variables of landscape heterogeneity at the scale of 1 km radius from the centroid of each farm: SNA (amount of total Semi-Natural Area, strongly correlated with Forest area; Pearson r = 0.945, P <0.001), ED (Edge Density), PD (patch diversity), PE (patch Evenness), OC (Amount of olive cover), PR (Patch Richness), LPS (Largest patch size), PA (mean Patch Area), NND (Distance to Nearest Neighbor patch of same type) and SDI (Shape Distribution Index). They were computed using the most recent and accurate land-use cartography of the region (SIOSE, http://www.siose.es), QGIS v.2.14 (QGIS Development Team, 2018) and FRAGSTATS software (McGarigal et al., 2012). Because some of them were highly correlated (see correlogram; Figure A.1, in Appendix A), we eventually selected 6 landscape variables that are the most commonly used and represented very well all the information: SNA, ED, PD, OC, PR, and NND.

### 2.3 Bird surveys

Birds were surveyed monthly from March 2016 to April 2017 (except July and August, 10 rounds) in six or ten permanent plots per farm (depending on farm size) separated by ca. 200m. Inside each farm, two (in small farms) or four (in big farms) of the plots were sited in patches of semi-natural habitat (non-crop plots hereafter), and the rest of the plots were inside the matrix of the olive plantation (crop plots hereafter). This, allowed to sample communities at the farm scale, capturing farm heterogeneity, and also enabled comparisons between plot types (habitat or microhabitat effects). In these plots, experienced ornithologists annotated the birds listened or seen for five minutes, within and outside a circle of 50m radius from the center of the plot. Censuses were conducted within three hours after sunrise. For this study, only insectivorous birds were accounted. Species were classified as insectivorous based on expert knowledge, and validated using the functional trait database by Storchová & Hořák (2018) and Wilman et al., (2014). Because forest birds might have a greater impact on pests (they forage more specifically on trees, where pests might be more vulnerable to birds), this same database was also used to classify birds as forest or open habitat species.

### 2.4 Pest abundance

We monitored the abundance of olive moth, by using funnel traps with pheromone z-7-tetradecen-1-ol that attracts adult males and an insecticide pill. Monitoring was conducted from April to July 2017, with monthly rounds to count trapped moths. We also monitored the abundance of adult olive flies in the 9 organic olive farms, from July to November 2016. For that, we set McPhail traps with the attractant ammonium bisulfate ((NH4) HSO4) diluted in water (4%). In every olive farm, we set a total of 6 (in small farms, < 25 ha) or 10 (in big farms, >100 ha) traps, hanging from olive trees widespread throughout the whole farm. McPhail traps were monthly checked and refilled with liquid.

### 2.5 Bird exclusion experiment

In this study, we measured the potential of insectivorous birds to control olive tree pests. We designed an experiment to preclude birds from accessing some branches. The damages observed in excluded branches were compared with those detected in control branches (parallel, close and similar to the experimental branch) that were not excluded from birds (e.g., Maas, Clough, & Tscharntke, 2013 for a similar approach). The aim of this experiment was to test the possible top-down effect of birds on olive tree damage, through arthropod/pest control. In each farm, 10 trees were selected, 5 non-consecutive trees in the first row adjacent to a patch of semi-natural habitat and 5 within the olive tree matrix (far from a patch of semi-natural habitat), ca. 100-120m away from the reference semi-natural patch. Bird mist-netting data from these two orchard positions clearly show that insectivorous bird activity is notably higher in the non-crop patches (authors’ unpublished data). Each selected tree had 4 excluded branches, and its corresponding control branches nearby. To exclude branches from birds, we used a cylinder of 80 cm long and 25 cm diameter made by 1cm^2^ pore plastic mesh. The ends were closed using 2mm^2^ pore plastic mesh (Figure A.2, in Appendix A). In total, we set 360 exclusions in 90 trees at the very beginning of March and were kept throughout the year. We conducted three check rounds, measuring four types of damages to leaves, flowers and fruits. Specifically, we sampled: i) leaf damage produced by leaf miner larvae of the olive moth and other phytophagous insects at the end of March; ii) leaf and flower damage by anthophagous larvae of the olive moth and other phyllophagous insects at the end of May; iii) leaf and fruit damage, by the moth, the fly and other phyllophagous insects at the end of October-beginning of November.

### 2.6 Artificial prey experiment

We employed plasticine models to compare the potential of avian predation pressure on two olive pest species across our landscape complexity gradient. This is a methodology commonly used in field experiments to assess and compare attack rates by specific predator guilds towards specific preys (Howe, Lövei, & Nachman, 2009; Lövei & Ferrante, 2017). We mimicked larvae of two generations of olive moth (phyllophagous generation in March-April and anthophagous generation in May) and the larval stage of olive fly (in October-December), being all the stages considered as temporal replicates of the same treatment. In each round, we set 4 plasticine models per tree, in 5 trees near a patch of semi-natural vegetation and 5 trees inside the olive orchard matrix. Selected olive trees were separated from others by at least another tree, ca. 20-30 m. Dummies were fixed to the ground, leaves or flowers depending on the pest and generation we tried to mimic (Figure A.2 in Appendix A). We conducted a total of 6 rounds per site (2 rounds per pest and larval stage), and a total of 2160 plasticine models were used (240 per site). We detected and counted predation marks by birds (beak mark) and by other predators (mainly ants, many small bites) on the plasticine models. All dummies were checked after 7 ± 1 day of exposure (Mäntylä et al., 2008) (with the exception of the first round where we tried 15 days ± 2 days). This method has been criticized in some systems for being not realistic (Zou et al., 2017), therefore we firstly set trap cameras to successfully validate some predation events by birds on our plasticine models.

### 2.7 Statistical analyses

First, we sought for patterns linking the abundance and richness of insectivorous birds to landscape complexity. Because some vulnerable stages of the studied pests are found on the ground, we first accounted for overall insectivorous birds. However, we reran the analyses accounting only for forest insectivorous birds, since this guild might have a greater impact on olive pests (they can forage on trees). For that, we ran Bayesian models with insectivorous bird abundance/bird richness as response variables and the six landscape components (see above, e.g. amount of semi-natural area, edge density, etc) at the scale of 1 km radius, as explanatory variables. Farm ID entered the models as a random factor. Response variables were log transformed to run normal models. We employed multilevel mixed Bayesian models fitted through the MCMC (Markov Chain Monte Carlo) method using the *brms* package (Bürkner, 2017) in *R* (R Core Team 2019). Results were interpreted as the posterior probability of beta (slope) being positive/negative. This kind of model enables us to run hierarchical mixed models. Also, Bayesian models are more easy and straightforward to interpret, especially for designs with a small-moderate number of cases, allow probabilistic predictions, and inferences are conditional on the data (which means that are exact). We also analyzed bird abundance/richness variation (for insectivorous birds and forest insectivorous birds) between plot types (non-crop plots and crop plots) within each farm. For this, we fitted Bayesian models, with bird abundance/richness as response variable and plot type as explanatory factor. Farm ID was also introduced as blocking random factor. We also explored differences in species assemblages, by analyzing how species-specific probability of occurrence varied across our range of landscape heterogeneity measured at 1 km radius, using Hierarchical Modelling of Species Communities (HMSC) (Ovaskainen et al., 2017). A detailed description of methodology, results and discussion of species assemblages’ variation is shown in Appendix B.

Second, we tested how pest abundance varied with landscape heterogeneity and bird abundance/richness (for overall insectivores and forest insectivores, separately), using Bayesian mixed models. We ran one model for each explanatory variable (six landscape components, bird abundance and bird richness) and each response variable (pest species; olive moth and olive fly). Farm ID was introduced as a random factor and the six landscape variables and bird abundance/richness as fixed factors. We inferred the results by testing the hypothesis of the posterior beta parameter being higher or lower than 0.

Third, we ran the same type of multilevel Bayesian mixed models, using the *brms* package to look for patterns in the damage observed in excluded and control branches. The response variable, damage, was arcsine square root transformed since it was a proportion (observed damage in relation to number of leaves, flowers and fruits counted). The explanatory variables were treatment (excluded vs. control) and site (close to a semi-natural patch or in the olive tree matrix, away from the semi-natural patch). Farm and tree ID were introduced as random factors. Additionally, we explored interactive effects of landscape complexity and distance to semi-natural patch on observed damage, because we expected less effect of distance in complex landscapes. These interactions did not explain variance in observed damage and were removed from models.

Finally, we tested how observed attack rates in pest dummies varied in relation to landscape components, bird abundance, bird richness, size of focal semi-natural patch and distance to semi-natural patch (as factor: close-distant). Attack rates (attacked/exposed) were arcsine square root transformed and linear mixed Bayesian models fitted. Farm ID was included as random factor in every model.

For all the Bayesian models, we checked convergence through R^ (all equal to 1 or 1.01), normality of the residuals and stability of results (by visual inspection of chains). We also inspected models’ goodness of fit via plots confronting observed data with posterior data generated using model simulations (N=200 datasets simulated). For all models, we used uninformative diffuse priors and model specifications that rendered stable outputs (4 chains and 50000 iterations with the first 10000 being burned).

All the analyses were run using *R*, version 3.6.1 (R Core Team 2019).

## 3. Results

During surveys, 86 different species of insectivorous birds, belonging to 62 genera and 36 families were identified; 47 of them were forest insectivores, from 36 different genera (see Table A.1 in Appendix A for the complete list of species). A number of 4966 insectivorous birds was detected across the year inside the 50m radius plots, of which 3349 were forest insectivores. The most abundant species were the sardinian warbler (*Sylvia melanocephala*; 15.9% of the total), the eurasian blackcap (*Sylvia atricapilla*; 11.8%), the common chaffin (*Fringilla coelebs*; 10.9%) and the european robin (*Erithacus rubecula*; 10.5%). Among bird families typically considered important for pest control in tree crops, both Paridae (4 species and 11% of total abundance) and Picidae (3 species and 0.2% of total abundance) were scarce in olive orchards. Thus, *Parus major* was the most frequent (8%) followed by *Cyanistes caeruleus* (3%) while *Lophophanes cristatus* and *Periparus ater* were rather rare and, as *Picus viridis, Dendrocopos major* and *Jynx torquilla*, appeared only associated to semi-natural and riparian forest patches.

The abundance of insectivorous birds did not vary importantly between olive farms surrounded by landscapes of different complexity (see Table 1). Only edge density increased the abundance of birds (Mean beta estimated = 0.14; Prob β > 0 = 0.94). However, species richness increased considerably in plots with landscapes showing a higher patch diversity (Mean beta estimated = 0.29; Prob β > 0 = 0.96). Accordingly, landscapes dominated by olive orchards (Mean beta estimated = −0.06; Prob β < 0 = 0.91) and with low distance to nearest similar patch decreased bird richness (Mean beta estimated = −0.11; Prob β < 0 = 0.99 for). These results were reinforced for forest insectivores. Their abundance increased in plots within landscapes with high density of edges (Mean beta estimated = 0.15; Prob β > 0 = 0.98) and their species richness increased in those with more semi-natural area (Mean beta estimated = 0.07; Prob β > 0 = 0.91) and edge density (Mean beta estimated = 0.12; Prob β > 0 = 0.98). Within farms, insectivorous birds were also strongly associated to non-crop patches (Table A.2, in Appendix A), showing lower abundance in crop patches (Mean beta estimated = −0.31; Prob β > 0 = 0). This pattern was stronger when only forest insectivores were considered, decreasing their abundance (Mean beta estimated = −0.40; Prob β > 0 = 0) and species richness (Mean beta estimated = −0.09; Prob β > 0 = 0.05) in crop plots. Additionally, we detected clear patterns of insectivorous bird species turnover as a function of landscape heterogeneity (see Appendix B).

**Table 1.**
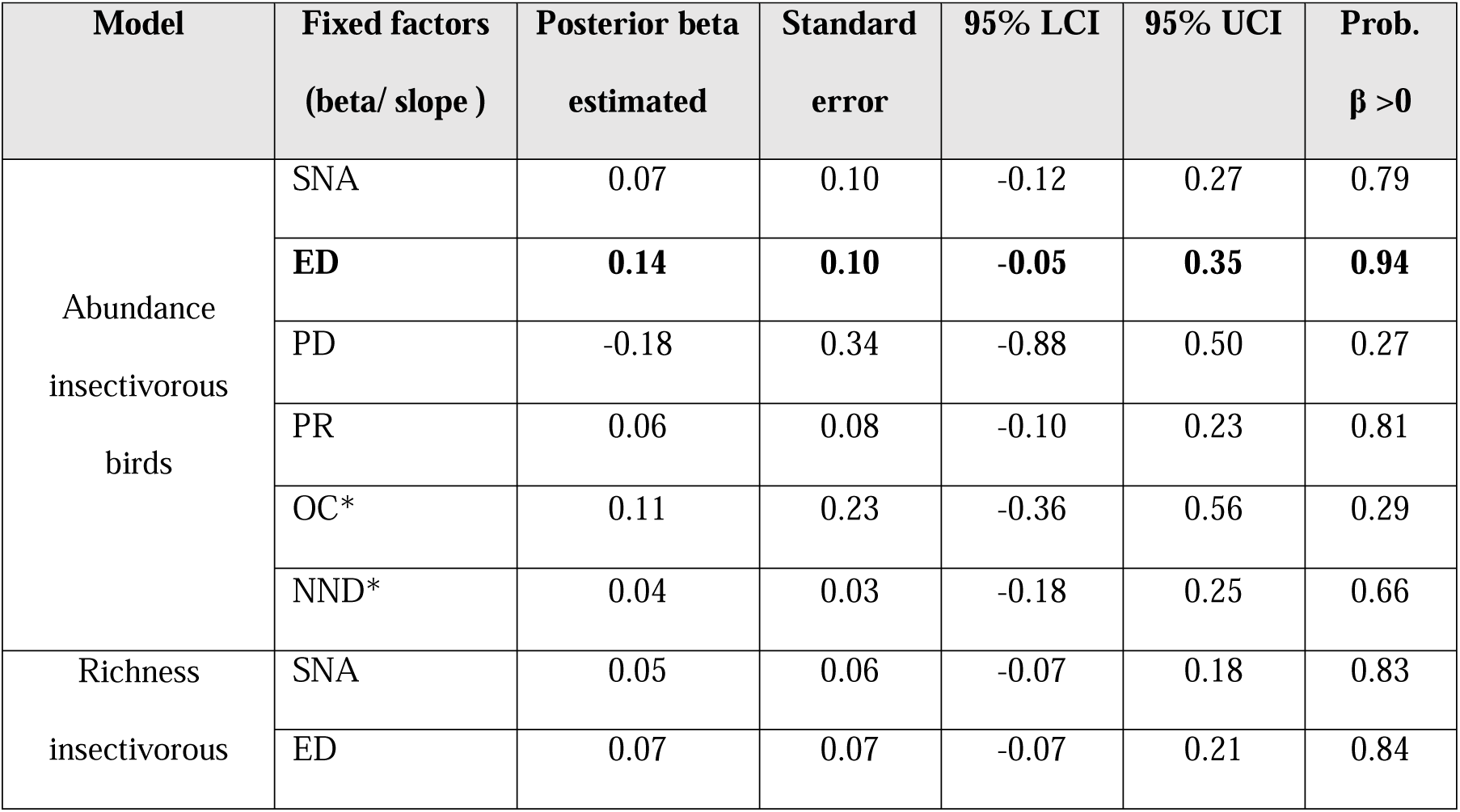
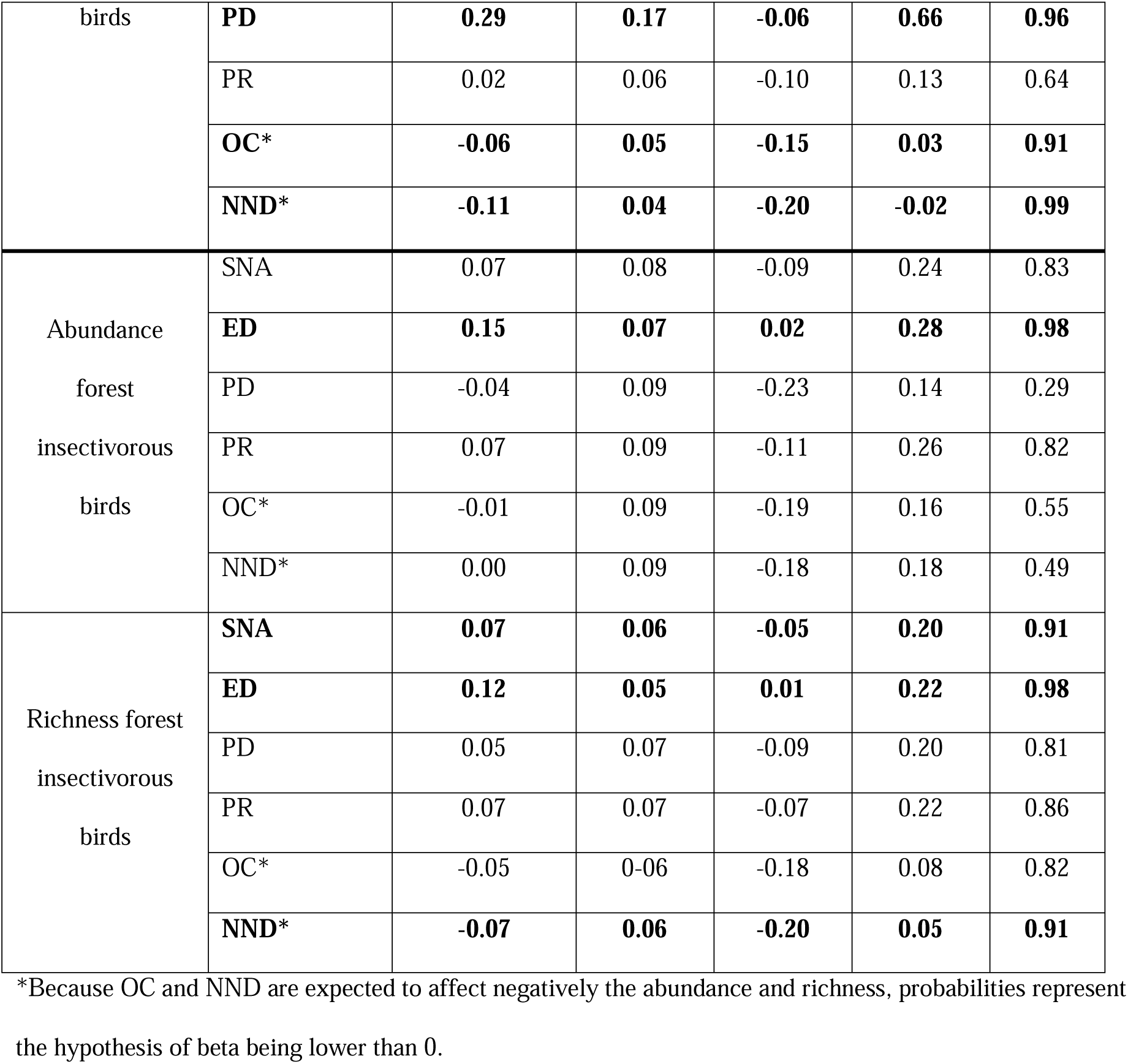
Results from Bayesian hierarchical models that show the estimated effect of landscape heterogeneity (scale of 1km radius) on the abundance/richness of insectivorous birds. The table displays the posterior mean, standard error, 95% credible intervals, and probability of beta being higher than 0. Results are in the log scale. In bold, models with a posterior probability over 90% of beta (slope) being positive/negative. SNA = Semi-Natural Area, ED = Edge Density, PD = Shannon diversity, PR = Patch richness, OC= Area of olive orchards, NND = Distance to Nearest Neighbor.

The abundance of pests captured with traps was notable in all olive orchards (ranging from 277±127 to 2216 ± 205 for the moth; and 21±17 to 625 ± 228 mean captures/trap ± 1SD for the fly). The abundance of the moth was correlated negatively with SNA, ED and PD (Posterior beta < 0 with a probability of 1) but the abundance of the fly did not show any remarkable pattern. Also, neither the moth nor the fly abundance were related to total insectivorous or forest insectivorous bird abundance or richness (see Table A.3 in Appendix A).

There were no differences between the control and excluded branches (see Table 2 and Fig. 1) regarding damage caused by the moth (Mean beta excluded branch= −0.01, Prob β > 0 = 0.25). However, excluded branches showed less damage for the fly (Mean beta excluded branch= −0.10, Prob β > 0 = 0), phyllophagous insects (Mean beta excluded branch= −0.05, Prob β > 0 = 0) and overall pests (Mean beta excluded branch= −0.04, Prob β > 0 = 0). Trees adjacent to semi-natural patches suffered less damage than trees sited in the olive matrix for all the type of damages measured (Mean beta for matrix trees = 0.04, Prob β > 0 = 1 for the moth; 0.05, Prob β > 0 = 0.98 for the fly; 0.06, Prob β > 0 = 1 for phyllophagous insects; and 0.05, Prob β > 0 = 1 for cumulated damage) (Table 2 and Fig. 1).

**Table 2.**
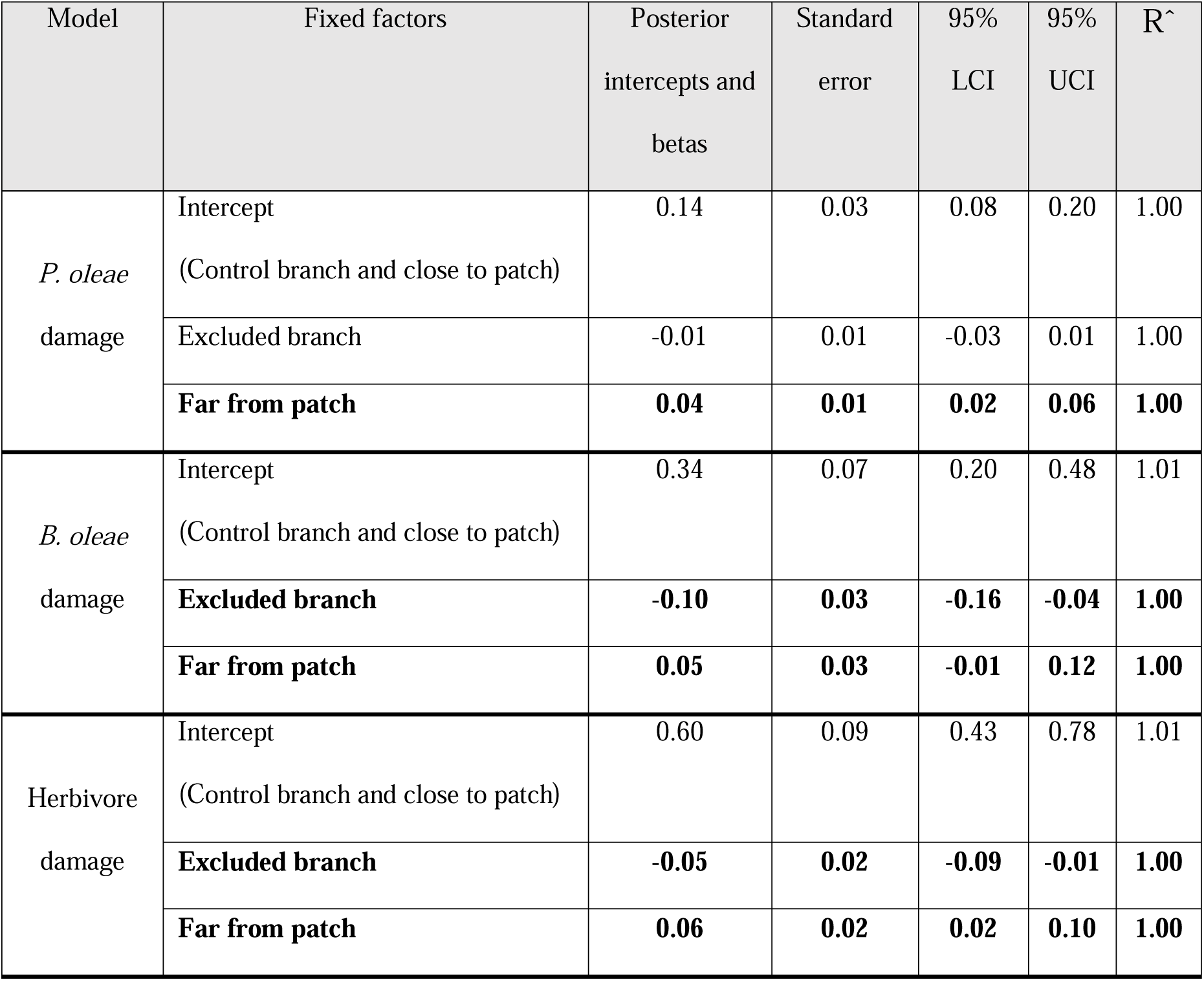
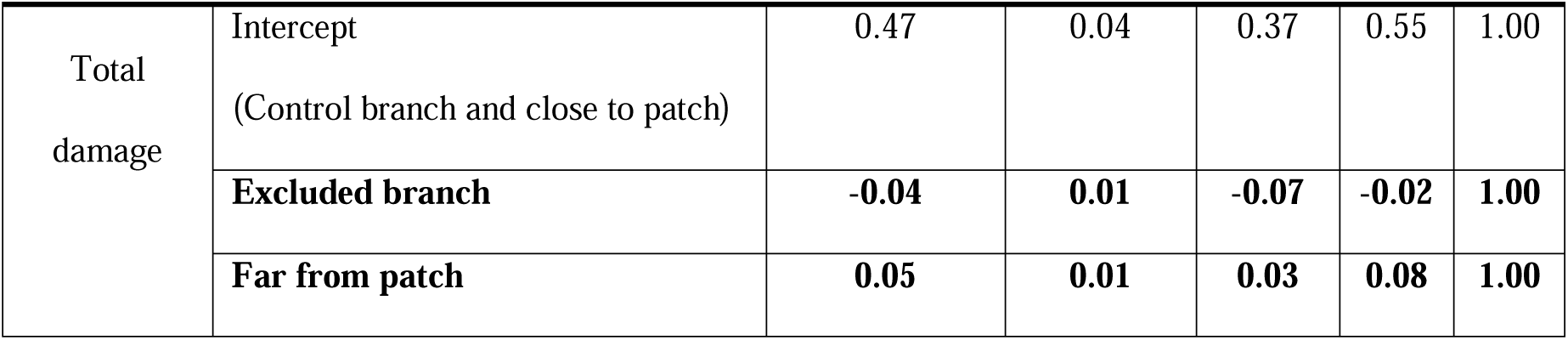
Results from Bayesian multilevel models that show the estimated effect of site (close or far from semi-natural patch) and treatment (excluded-control branch) on total damage observed to olive trees. The table displays the posterior mean, standard error, 95% credible interval, and R^ statistic for each parameter of models with a varying intercept by locality and tree. In bold, factor levels with a β parameter higher or lower than 0 (probability of 95%).

**Figure 1.**
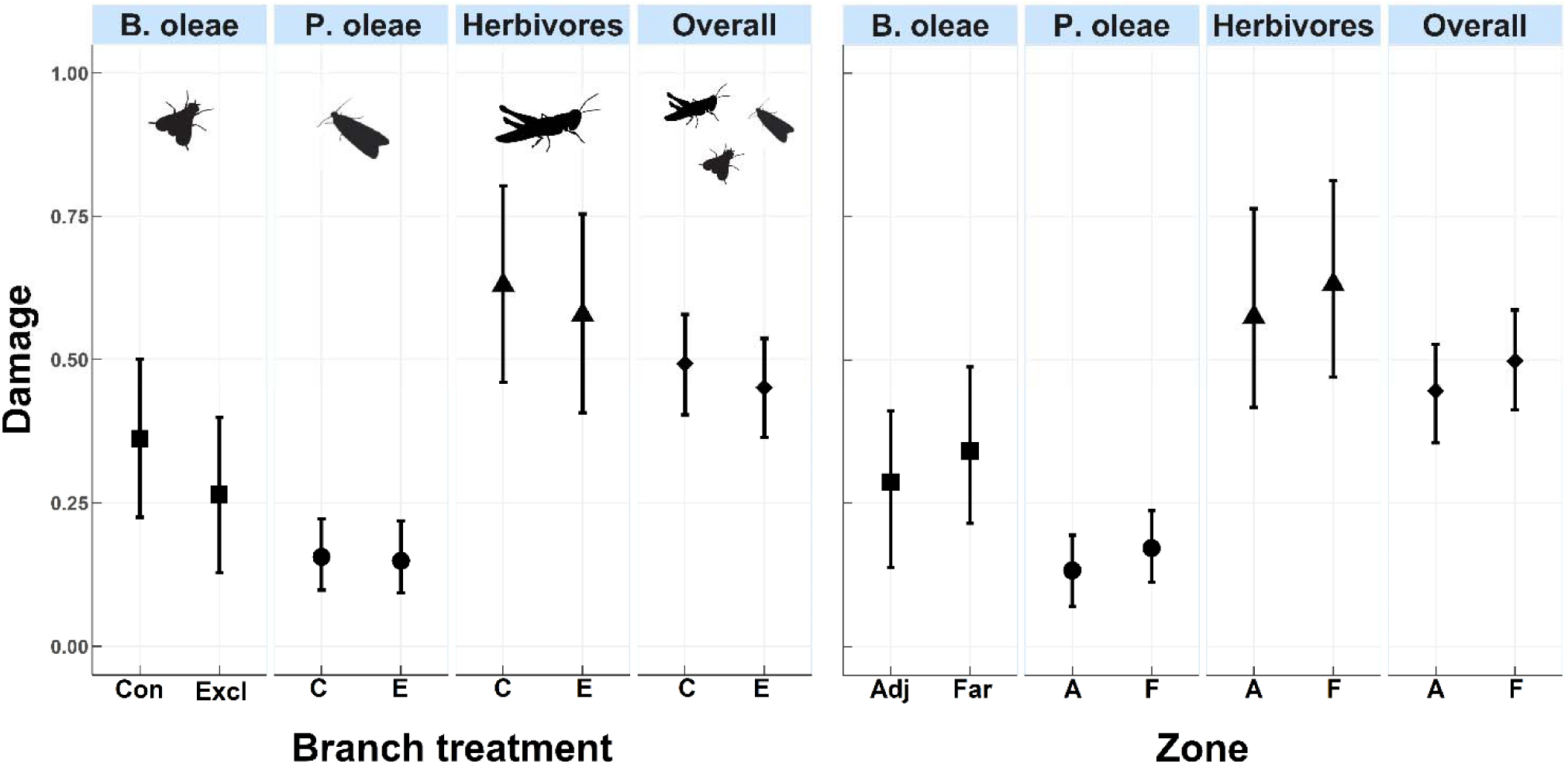
Back-transformed predicted damage (proportion) produced by *P. oleae, B. oleae*, Herbivores and Overall (cumulated) and its variation across treatments (Control Vs. Excluded branch) and Zone (Adjacent to semi-natural patch =Adj. Vs. olive orchard matrix, far from semi-natural patches= Far). Solid symbols show predicted posterior mean and whiskers 95% credible intervals.

The observed attack rates in plasticine models were extremely low, N = 144 bird attacks (less than 0.8% d^−1^) and N = 344 total attacks (by birds plus others, slightly over 1.9 % d^−1^) (Fig. 2). There were no meaningful patterns explained either by landscape characteristics or distance to semi-natural patches, which probably means that the low attack rates were insufficient to show reliable patterns across landscapes or distance to seminatural patches.

**Figure 2.**
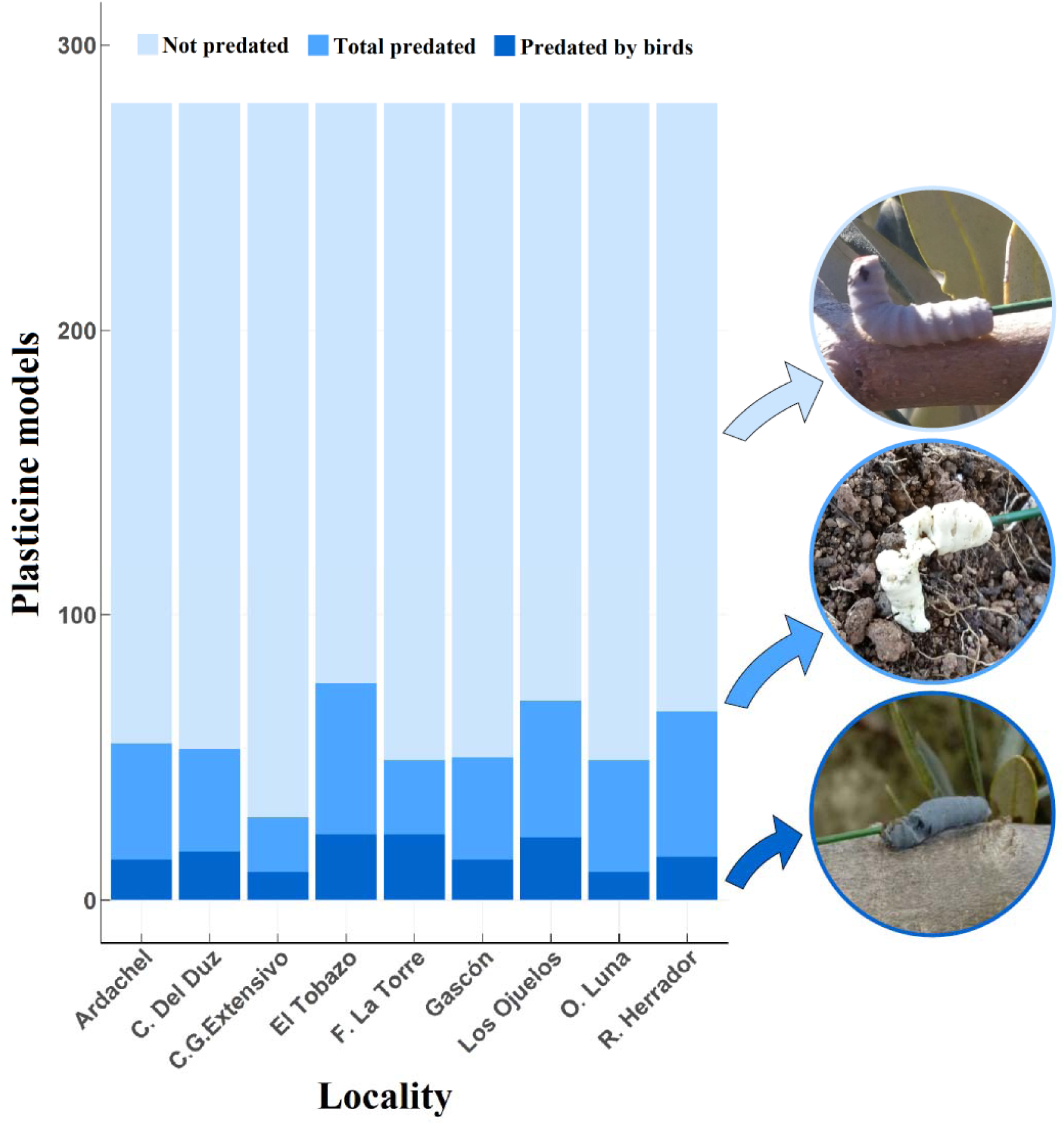
Observed number of plasticine models attacked by birds (dark blue), by other animals, mainly ants (clear blue) and not attacked (light blue).

## 4. Discussion

The results obtained supported our first prediction, since heterogeneous landscapes and non-crop habitats tended to host a higher abundance, richness, and a particular assemblage of insectivorous birds (particularly benefitting forest insectivores). However, excluded branches did not suffer more damage by pests, and attack rates to plasticine models were too scarce to show reliable patterns across landscapes and farms. Therefore, none of our two last predictions were supported by the results, suggesting that the pest control ecosystem service provided by insectivorous birds in olive orchards might be in a worrying situation.

### 4.1 Landscape effects on insectivorous birds

Despite the relatively low number of olive orchards considered, we did find a positive trend in insectivorous bird abundance and richness driven by some components of landscape heterogeneity. Interestingly, such differences were stronger for abundance and richness of forest insectivores. Consistently with other studies (e.g. García, Miñarro & Martínez-Sastre, 2018; Karp et al., 2013), more species of forest insectivores were found in landscapes with a higher proportion of semi-natural habitats. Results also showed important differences at the farm scale, with non-crop plots supporting a more abundant and rich community of insectivores (particularly forest insectivores) than crop plots. The underlying reason for these findings might be that complex landscapes offer more variability of niches, resources and higher food availability, being able to support a wider community of these insectivorous birds. This is further supported by our finding of important species turnover across our landscape gradients (Appendix B).

### 4.2 Effectiveness of pest control by insectivorous birds

The experiment with bird exclusions supported the lack of effective pest control by current populations of insectivorous birds in olive orchards. While other studies conducted in orchards and agroforests have found more damage in branches/trees excluded to birds (García, Miñarro & Martínez-Sastre, 2018; Maas, Clough, & Tscharntke, 2013), we found similar or lower damage in excluded branches compared to control branches across all our study sites. For the fly and other phyllophagous insects, we found more damage in control branches. These results might be due to intraguild predation, with birds predating on other insect predators such as ants or spiders in control branches, and thus releasing pests from these mesopredators (Maas et a. 2016). In any case, birds did not reduce pest damage. Interestingly, trees close to semi-natural patches showed less damage than trees located inside the olive tree matrix. This decrease in damage was not affected by branch treatment (excluded or control), which suggests that some small arthropods associated to semi-natural patches are effective natural enemies against these two olive pests. Their predation pressure might influence pest preference for inner trees or directly reduce pest populations in trees close to semi-natural patches. Some studies have pointed out the key role of arthropod natural enemies such as *Anthocoris nemoralis* (Fabricius) (Paredes et al., 2019) or other ground arthropods (Dinis et al., 2016) in olive orchards. Semi-natural patches and ground cover seem to have a vital function, providing habitat and alternative prey for them (Álvarez et al., 2019; Paredes et al., 2019). This result endorses the positive effect of natural fragments on pest control in agroecosystems, and the higher predation observed in areas close to semi-natural patches/ecotones, already reported by other authors in different systems (Barbaro, Giffard, Charbonnier, van Halder, & Brockerhoff, 2014; Maas et al., 2015; Milligan, Johnson, Garfinkel, Smith, & Njoroge, 2016).

The recorded attack rates to plasticine models did not follow any pattern associated to landscape features. The results obtained in this experiment agree with the previous experiment (i.e. branch exclusions) because attack rates by birds were extremely low. Despite that recorded predation is usually higher on real preys than on dummy (Lövei & Ferrante, 2017), observed numbers are well under attack rates needed for an effective pest control. Higher attack rates on dummy preys have been found in other agricultural systems such as: oil palm plantations (37% without and 53% with riparian forest fragments) (Gray & Lewis, 2014), apple plantations (63%, Martínez-Sastre et al. in press), cacao agroforest (2.9% d^−1^ attack rate by birds, and 6.5% d^−1^ including all arthropods) (Maas et al., 2015) or cotton fields (ca. 4% d^−1^) (A. G. Howe, Nachman, & Lövei, 2015). As noted by Lövei & Ferrante (2017), relative low densities of natural preys can increase attack rates to dummy preys. Some authors have shown this effect by detecting increased attack rates in more disturbed areas (rural areas vs. open and closed canopy forests) (Posa, Sodhi, & Koh, 2007). Large numbers of these pests, added to a relatively low abundance of insectivorous birds using olive orchards could explain the negligible attack rates found in our study (see next section). Some authors have found a correlation between attack rates (to plasticine models) and the abundance of insectivorous birds surveyed with mist-nets (Maas et al., 2015). This method might be better than conventional surveys to address the birds that are really entering in the olive tree matrix, in contrast to birds that use the semi-natural patches but are rarely found inside the olive orchards.

### 4.3 Functional dilution hypothesis

We recorded a high number of adult pests with traps (mean of 1372/trap for olive moth, and 134/trap for olive fly), and the observed damage was high as well (ca. 15% damaged by olive moth; ca. 30% by olive fly; and ca. 60% of leaves damaged by phyllophagous insects), evidencing a high density of the moth, the fly, and phyllophagous insects in the studied farms. However, despite this temporal high availability of prey, insectivorous birds seem to make a limited use of olive orchards and crop plots inside farms, as to be effective controlling these pests. This overall limited use of olive orchards would indicate low attractiveness of the olive matrix for avian insectivore guilds, especially forest insectivores. This is supported by the fact that insectivorous bird assemblages clearly augmented in non-crop patches within farm compared to the crop matrix, this being even more marked for forest insectivores. Some other studies have also shown that the richness and abundance of insectivorous-frugivorous birds decline (at least in autumn-winter) in olive groves compared to semi-natural wild olive scrublands and other reference woodlands (Alcántara et al., 1997a; Rey, 1993). These declines in avian insectivore may suppose well above one order of magnitude in terms of energetic requirement by hectare of the whole insectivore guild between these habitats (Alcántara et al. 1997b). Forest insectivores have also been shown to decline in olive orchards during the reproductive season as a function of agriculture intensification and/or landscape simplification (Castro-Caro, Barrio, & Tortosa, 2015; Morgado et al. 2020). Such limited attractiveness of the olive tree matrix may be related with the current suboptimal (structural and feeding) conditions of this habitat where the management originates regular distribution of trees, absence of nesting cavities on tree trunks, short and reduced tree canopies, and lack of permanent scrub and/or herbaceous layers and their associated food resources.

On the other hand, the observed mismatch between prey-predator abundance might be partially caused by relatively scarce and small semi-natural areas at the olive farm and landscape scales. Thus, there are little resources (e.g. alternative preys or nesting sites) to host a bird community large enough to exert an important insectivory pressure. Moreover, the ratio natural to crop area needed in Mediterranean agroecosystems to effectively sustain insectivorous bird populations should be higher than this ratio in other places (e.g. tropical areas), because semi-natural patches have typically low productivity in Mediterranean environments (as a consequence of the severe summer drought) compared to tropical and non-Mediterranean temperate ones (Imhoff & Bounoua, 2006). This likely prevents the permanent presence of large enough populations and rich assemblages of insectivorous birds within olive orchards.

Therefore, it is likely that avian insectivore density in the tree matrix of olive farms at any landscape was under the threshold levels needed for an effective pest control. We argue, thus, for a functional dilution hypothesis to explain the non-effective pest control by birds in olive orchards. This hypothesis states that avian predation pressure and control over pests is diluted at the landscape level due to low numbers of predators and natural habitat, compared to the abundance of pests and the extent of crop area. Similar findings were reported by McConkey & Drake (2006), where the authors detected the loss of an ecosystem service (seed dispersal) provided by bats, well before important bat population decline.

To our knowledge, this is the first study to assess pest control by birds in olive orchards, which hinders further comparison of our results. However, Rey Benayas et al., (2017) set an experiment in several Mediterranean woody crops, including olive orchards, using bird nest boxes and alive sentinel preys. The nest boxes set in olive orchards were not occupied throughout the four-year study, and therefore, no attack rates were available for this system. Lack of occupation of nest boxes in this system is congruent with our results reporting a worrying situation for insectivorous birds and their contribution to pest control in olive orchards. Nonetheless, we cannot discard that our results on pest control by birds might be reflecting a relatively low bird appetence for the two pest species studied (Tscharntke et al., 2016). Unfortunately, there is a notable gap in studies analyzing bird insectivore diet in olive groves in relation to its major pests, whereby there is no information of reference. As done in other agroecosystems (e.g., Karp et al. 2013 in coffee plantations or Mangan et al. 2018 in apple orchards), studies analyzing bird diet with molecular tools (e.g. DNA analyses form faecal sampling) are needed to elucidate how frequently insectivorous birds feed on olive pests.

## Conclusion

This study suggests that insectivorous birds are unable to provide an effective biological control service for the two main olive tree pests in Andalusia (Spain). Our results might be showing the serious consequences of current low insectivorous bird densities on the pest control ecosystem service carried out in olive orchards. Since alarming bird declines have been reported in North America (Rosenberg et al., 2019) and Europe in the last decades (Bowler, Heldbjerg, Fox, de Jong, & Böhning-Gaese, 2019; Inger et al., 2015), avian insectivory function in olive orchards is expected to be aggravated in the olive farms in the near future, since olive matrix is clearly suboptimal for avian insectivores, and suboptimal habitats become empty earlier that optimal ones. According to our results, urgent actions should be taken to reverse this situation, such as policy changes aimed to the recuperation of landscape heterogeneity and semi-natural patches in landscapes homogenized by monoculture, and active restoration of bird assemblages (e.g. setting bird nest boxes and feeders).

## Author’s contributions

PJR, DG and CMN conceived the ideas and sampling design of this study. TS and PJR conducted the landscape analysis. CMN, RT, JLM, AJM, and PJR field experiments and carried out pest abundance and damage surveys and RT, JLM, and PJR the bird censuses. CMN led the analysis and wrote the first draft of the manuscript with feedback from PJR and DG. All authors contributed significantly to the final version.

## Acknowledgements

We thank the farmers who collaborated with us, granting access to their properties. We also thank Antonio J. Pérez, Gemma Calvo, Maria José Navarro, Francisco Camacho, Enrique Muñoz and Sandra Lendínez for helping with fieldwork. Finally, we thank Francisco Valera, José E. Gutiérrez and Carlos Ruiz for logistic support. This work is part of the projects AGRABIES (CGL2015□68963□C2, MINECO, Gobierno de España and FEDER) and LIFE OLIVARES VIVOS (LIFE14 NAT/ES/001094, European Commission). CMN was granted a predoctoral fellowship (BES-2016-078736). Authors have no conflict of interest.

## Data deposition information

Should the manuscript be accepted, data will be uploaded to the Mendeley Data repository.

## Ethics declarations

### Informed consent

Informed consent was obtained from all individual participants included in the study.

