## Appendix A for "Landscape drivers and effectiveness of pest control by insectivorous birds in a landscape-dominant woody crop"

^1^ Dept. Biología Animal, Biología Vegetal y Ecología, Universidad de Jaén. E-23071 Jaén, Spain.

^2^ Instituto Interuniversitario del Sistema Tierra de Andalucía, Universidad de Jaén, E-23071

Jaén, Spain.

^3^ Centro de Estudios Avanzados en Ciencias de la Tierra, Energía y Medio Ambiente. Universidad de Jaén, E-23071 Jaén, Spain.

^4^ Departamento de Biología de Organismos y Sistemas, Universidad de Oviedo, y Unidad Mixta de Investigación en Biodiversidad (CSIC-Uo-PA), C/Catedrático Rodrigo

Uría s/n, E-33006, Oviedo, Asturias, Spain

^5^ Estación Experimental de Zonas Áridas. Almería. Spain

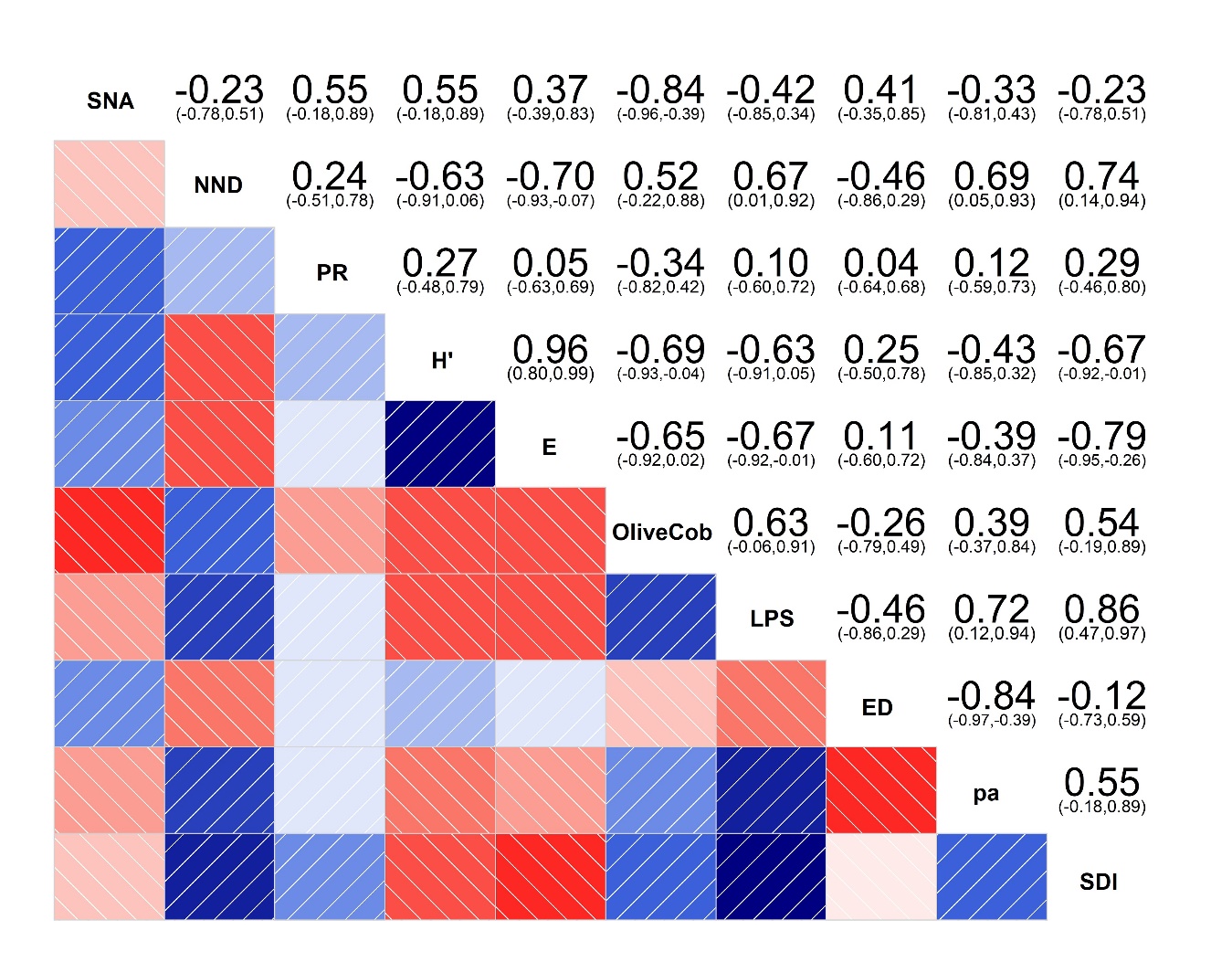

**Figure A1:** Correlogram showing how landscape variables correlate with each other. SNA = Semi-natural area, NND = Distante to Nearest Neighbour, PR= Patch Richness, H’ =Shannon Diversity of patches, E = Evenness of patches, OliveCob = Olive area, LPS = Largest Patch Size, ED = Edge Density, pa = Mean Patch Size, and SDI = Shape Distribution Index.

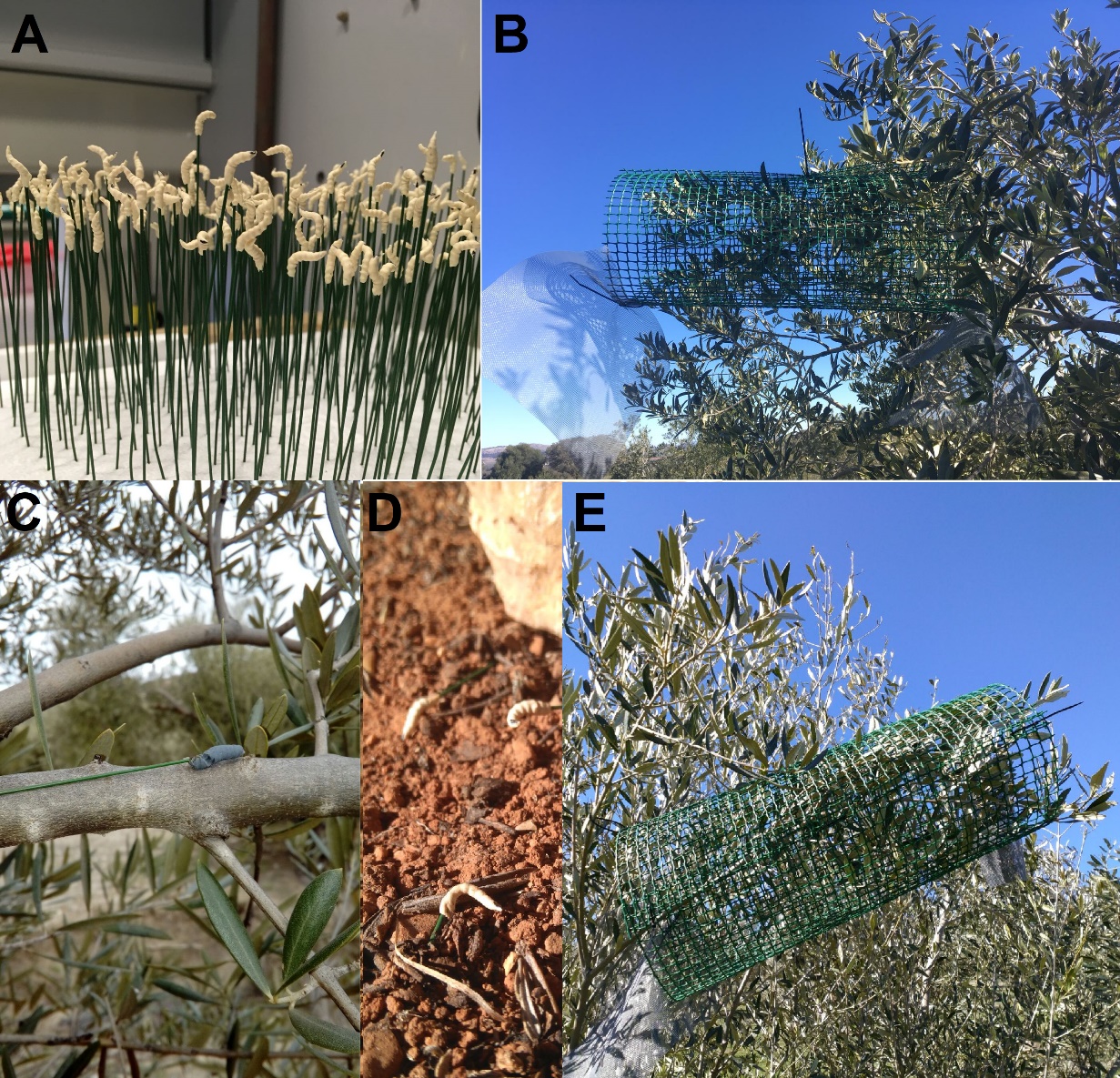

**Figure A2:** Pictures of plasticine models of *B. oleae* prepared in the lab (A), *P. oleae* phyllophagous stage with predation marks (C) and *B. oleae* in the field; displayed various together for a preliminary experiment (camera trap) (D). Pictures B and E show olive branches with exclusions (still with the ends open).

**Table A1:** Insectivorous bird species detected during the surveys, percentage of insectivory in their diets and the habitat they live in (F= Forestal; 1=Yes, 0=No). According to Wilman, H. et al. (2014), and Storchova et al. (2018). Note that some species with low percentage of insectivory in this database (e.g. Corvus corax or Lanius excubitor), can be locally important insectivorous (expert knowledge).

| **Species** | **Insectivory (%)** | **F** | **Species** | **Insectivory (%)** | **F** |
| --- | --- | --- | --- | --- | --- |
| Acrocephalus schoenobaenus | 70 | 0 | Hirundo rustica | 80 | 0 |
| Acrocephalus scirpaceus | 70 | 0 | Iduna pallida | 80 | 1 |
| Aegithalos caudatus | 60 | 1 | Jynx torquilla | 100 | 1 |
| Alauda arvensis | 40 | 0 | Lanius excubitor | 10 | 1 |
| Anthus campestris | 70 | 0 | Lanius senator | 80 | 1 |
| Anthus pratensis | 80 | 0 | Lophophanes cristatus | 60 | 1 |
| Anthus spinoletta | 70 | 0 | Lullula arborea | 50 | 0 |
| Anthus trivialis | 60 | 1 | Luscinia megarhynchos | 70 | 1 |
| Apus apus | 100 | 0 | Melanocorypha calandra | 50 | 0 |
| Apus pallidus | 100 | 0 | Merops apiaster | 100 | 0 |
| Bubulcus ibis | 60 | 0 | Monticola solitarius | 40 | 0 |
| Burhinus oedicnemus | 50 | 0 | Motacilla alba | 100 | 0 |
| Calandrella brachydactyla | 60 | 0 | Motacilla cinerea | 100 | 0 |
| Cecropis daurica | 100 | 0 | Muscicapa striata | 80 | 1 |
| Certhia brachydactyla | 70 | 1 | Oenanthe hispanica | 60 | 0 |
| Cettia cetti | 90 | 0 | Oriolus oriolus | 30 | 1 |
| Cisticola juncidis | 80 | 0 | Otis tarda | 40 | 0 |
| Clamator glandarius | 90 | 1 | Otus scops | 80 | 1 |
| Coracias garrulus | 90 | 0 | Parus major | 40 | 1 |
| Corvus corax | 10 | 1 | Passer hispaniolensis | 20 | 0 |
| Corvus corone | 30 | 1 | Periparus ater | 40 | 1 |
| Corvus monedula | 30 | 1 | Pernis apivorus | 80 | 1 |
| Coturnix coturnix | 20 | 0 | Phoenicurus ochruros | 60 | 0 |
| Cuculus canorus | 90 | 1 | Phoenicurus phoenicurus | 80 | 1 |
| Cyanistes caeruleus | 50 | 1 | Phylloscopus bonelli | 90 | 1 |
| Cyanopica cyanus | 80 | 1 | Phylloscopus collybita | 80 | 1 |
| Delichon urbicum | 100 | 0 | Phylloscopus trochilus | 80 | 1 |
| Dendrocopos major | 50 | 1 | Pica pica | 20 | 1 |
| Emberiza calandra | 30 | 0 | Picus sharpei | 90 | 1 |
| Emberiza cia | 20 | 1 | Ptyonoprogne rupestris | 100 | 0 |
| Emberiza cirlus | 30 | 1 | Regulus ignicapilla | 100 | 1 |
| Erithacus rubecula | 40 | 1 | Saxicola rubetra | 70 | 0 |
| Ficedula hypoleuca | 100 | 1 | Saxicola torquatus | 70 | 0 |
| Fringilla coelebs | 60 | 1 | Sitta europaea | 70 | 1 |
| Galerida cristata | 40 | 0 | Sturnus unicolor | 20 | 0 |
| Galerida theklae | 60 | 0 | Sturnus vulgaris | 20 | 0 |
| Garrulus glandarius | 40 | 1 | Sylvia atricapilla | 50 | 1 |
| Hippolais polyglotta | 80 | 1 | Sylvia borin | 50 | 1 |
| **Species** | **Insectivory (%)** | **F** |  |  |  |
| Sylvia cantillans | 60 | 0 |  |  |  |
| Sylvia hortensis | 70 | 1 |  |  |  |
| Sylvia melanocephala | 50 | 0 |  |  |  |
| Sylvia undata | 70 | 0 |  |  |  |
| Tachymarptis melba | 100 | 0 |  |  |  |
| Troglodytes troglodytes | 60 | 1 |  |  |  |
| Turdus iliacus | 40 | 1 |  |  |  |
| Turdus merula | 50 | 1 |  |  |  |
| Turdus philomelos | 40 | 1 |  |  |  |
| Turdus viscivorus | 40 | 1 |  |  |  |
| Upupa epops | 80 | 1 |  |  |  |

**Table A2.** Results from Bayesian hierarchical models that show the estimated effect of plot type (non-crop plot or crop plot) on the abundance/richness of insectivorous birds and forest insectivorous birds. The table displays the posterior estimate, standard error, 95% credible intervals, and probability of beta being higher than 0. Results are in the log scale. In bold, estimates with a posterior probability over 90% of beta (slope) being negative.

| **Model** | **Fixed factors**  **(beta/ slope )** | **Estimate** | **Standard error** | **95% LCI** | **95% UCI** | **Prob.**  **β >0** |
| --- | --- | --- | --- | --- | --- | --- |
| Abundance insectivorous birds | Non-crop plots  (intercept) | 4.28 | 0.11 | 4.06 | 4.50 |  |
|  | **Crop plots** | **-0.31** | **0.07** | **-0.44** | **-0.18** | **0** |
| Richness insectivorous birds | Non-crop plots  (intercept) | 3.16 | 0.07 | 3.02 | 3.30 |  |
|  | Crop plots | -0.00 | 0.03 | -0.07 | 0.06 | 0.48 |
| Abundance forest insectivorous birds | Non-crop plots  (intercept) | 3.91 | 0.10 | 3.70 | 4.11 |  |
|  | **Crop plots** | **-0.40** | **0.09** | **-0.56** | **-0.23** | **0** |
| Richness forest insectivorous birds | Non-crop plots  (intercept) | 2.21 | 0.09 | 2.03 | 2.39 |  |
|  | **Crop plots** | **-0.09** | **0.06** | **-0.21** | **0.02** | **0.05** |

**Table A3.** Results from Bayesian hierarchical models that show the estimated effect of insectivorous bird abundance and richness on the abundance of *P. oleae* and *B. oleae*. The table displays the estimates, standard error, 95% credible intervals, and probability of beta being lower than 0 (i.e. higher abundance of insectivores are related to lower abundance of pest).

| **Model** | **Fixed factors**  **(beta/ slope )** | **Estimate** | **Standard error** | **95% LCI** | **95% UCI** | **Prob.**  **β < 0** |
| --- | --- | --- | --- | --- | --- | --- |
| Abundance  *P. oleae* | Abundance insectivorous | -0.00 | 0.00 | -0.01 | 0.00 | 0.85 |
|  | Richness insectivorous | -0.01 | 0.02 | -0.05 | 0.02 | 0.77 |
|  | Abundance forest insectivorous | -0.00 | 0.00 | -0.01 | 0.00 | 0.82 |
|  | Richness forest insectivorous | -0.01 | 0.03 | -0.07 | 0.04 | 0.71 |
| Abundance  *B. oleae* | Abundance insectivorous | -0.00 | 0.01 | -0.01 | 0.01 | 0.71 |
|  | Richness insectivorous | -0.00 | 0.04 | -0.08 | 0.07 | 0.53 |
|  | Abundance forest insectivorous | -0.00 | 0.01 | -0.02 | 0.01 | 0.69 |
|  | Richness forest insectivorous | 0.04 | 0.06 | -0.07 | 0.15 | 0.23 |
