## Appendix B for "Landscape drivers and effectiveness of pest control by insectivorous birds in a landscape-dominant woody crop"

^1^ Dept. Biología Animal, Biología Vegetal y Ecología, Universidad de Jaén. E-23071 Jaén, Spain.

^2^ Instituto Interuniversitario del Sistema Tierra de Andalucía, Universidad de Jaén, E-23071

Jaén, Spain.

^3^ Centro de Estudios Avanzados en Ciencias de la Tierra, Energía y Medio Ambiente. Universidad de Jaén, E-23071 Jaén, Spain.

^4^ Departamento de Biología de Organismos y Sistemas, Universidad de Oviedo, y Unidad Mixta de Investigación en Biodiversidad (CSIC-Uo-PA), C/Catedrático Rodrigo

Uría s/n, E-33006, Oviedo, Asturias, Spain

^5^ Estación Experimental de Zonas Áridas. Almería. Spain

**How does probability of occurrence of specific species vary across landscape gradients?**

Using landscape metrics and bird species’ frequency data, we explored differences in species assemblages, by analyzing how species-specific probability of occurrence varied across our range of landscape heterogeneity measured at 1 km radius. For this analysis, we considered also bird species detected within the farm but outside the 50m radius plot; hence, in this case, we transformed abundance data to occurrence data and used Hierarchical Modelling of Species Communities (HMSC) (Ovaskainen et al., 2017). We used occurrence per plot as response variable and each landscape characteristic as explanatory variable. This Bayesian approach allows the specification of nested structures (e.g. N = 78 bird stations inside N = 9 farms) and identifies positive and negative responses of species to environmental gradients (i.e. “winners” and “losers”). We ran six models, one for each landscape feature, and used a Bernouilli (probit) distribution for presence-absence data. We removed from this analysis the species with 5 or less occurrences, because their models were not robust. Farm ID was introduced as a random level. Species that had a positive or negative posterior beta parameter (slope) with a probability equal or higher to 95% were plotted. This means that there is a 95% probability that the occurrence of a specific species covaries with landscape. We inspected the area under the receiver operating characteristics (AUROC) to check the degree of separability of each model for each species. We ran these models using the *Hmsc* (Tikhonov et al., 2019) package in *R*. We checked convergence through R^ (all equal to 1 or 1.01), normality of the residuals and stability of results (by visual inspection of chains). We also inspected models’ goodness of fit via plots confronting observed data with posterior data generated using model simulations (N=200 datasets simulated). For all models, we used uninformative diffuse priors and model specifications that rendered stable outputs (4 chains and 50000 iterations with the first 10000 being burned). All the analyses were run using *R*, version 3.6.1 (R Core Team 2019).

Using this methodology, we found important species turnover across our landscape gradients (see Fig. 1 in this Appendix). Olive farms located in landscapes characterized by a higher landscape complexity (i.e. higher amount of SNA, ED, patch richness or patch diversity), hosted more species typically associated to forests, such as *F. coelebs*, *P. bonelli*, *R. ignicapilla*, *C.* *cyanus*, *S. atricapilla* or *G. glandarius*, while many other forest insectivores did not show any significant trend (Paridae, Picidae), probably because they were rare in all landscapes. This suggests that olive farms with more semi-natural patches and located in complex landscapes might have a higher value for conservation of some forest insectivores and might benefit from the ecosystem services they provide. Congruent with this, olive orchards located in simple landscapes, showed a decline in insectivorous species richness (see main text). A recent study with breeding birds in olive orchards in Portugal showed an increase of generalist and open land species and a decrease of forest species with management intensification and landscape homogeneity (see Morgado et al., 2020). In any case, generalist and open land bird species are well adapted to agricultural areas and might be of vital importance for the ecosystem services provided in these simplified environments.

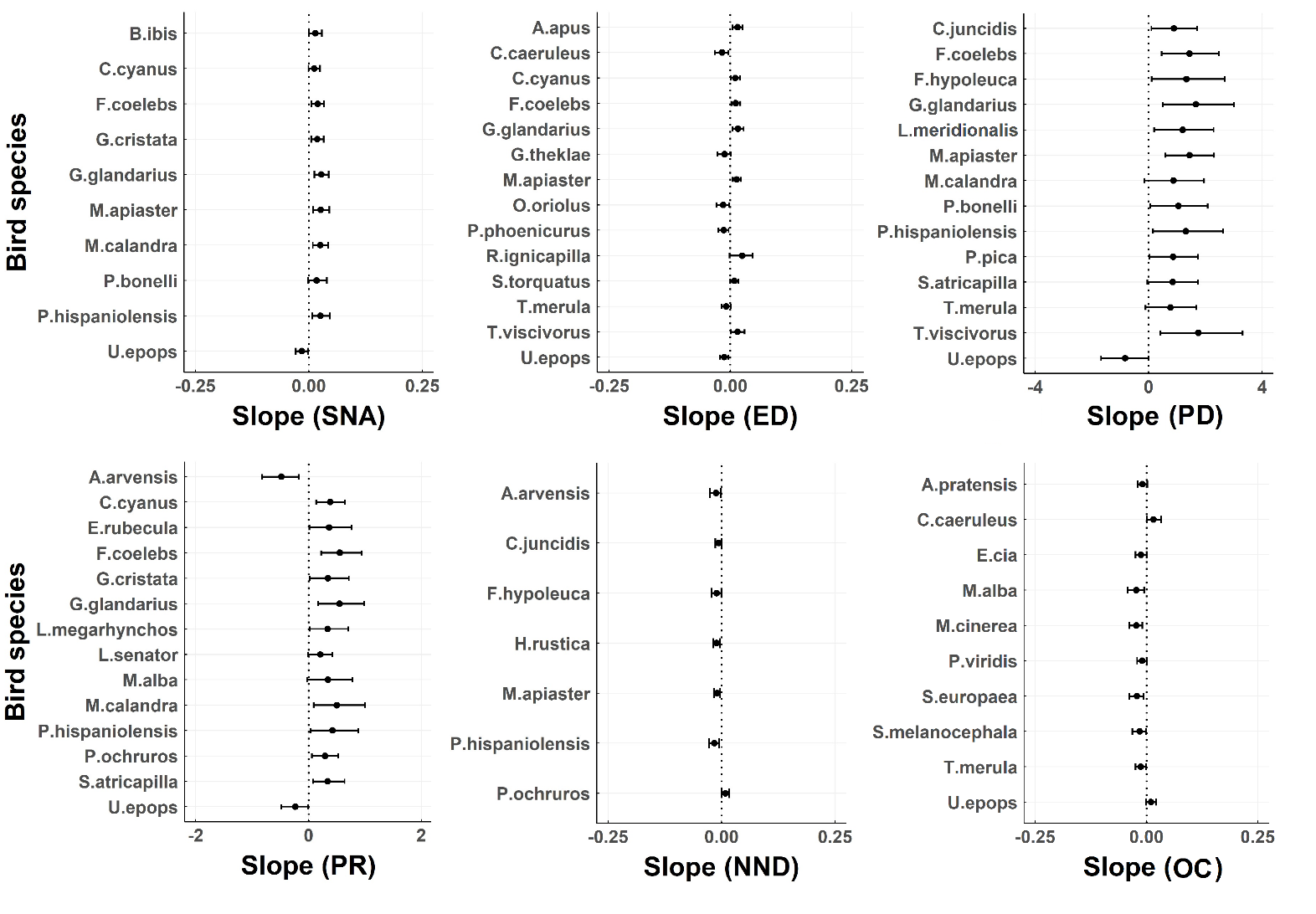

Figure 1. Results showing covariation between the abundance of bird species and landscape heterogeneity (HMSC) at 1 km radius buffer scale. Only relations with a support equal or higher to 95% probability are displayed. Negative relationships are represented with bars to the left (negative slope), positive relationships (positive slope) to the right. SNA = Semi-Natural Area, ED = Edge Density, PD = Shannon diversity, PR = Patch richness, NND = Distance to Nearest Neighbor, OC = Area of olive orchards.

Tikhonov, G., Opedal, Ø., Abrego, N., Lehikoinen, A., & Ovaskainen, O. (n.d.). Joint species distribution modelling with HMSC-R. https://doi.org/10.1101/603217
